# Human myelin proteolipid protein structure and lipid bilayer stacking

**DOI:** 10.1101/2022.03.14.484199

**Authors:** Salla Ruskamo, Arne Raasakka, Jan Skov Pedersen, Anne Martel, Karel Škubník, Tamim Darwish, Lionel Porcar, Petri Kursula

**Affiliations:** Faculty of Biochemistry and Molecular Medicine & Biocenter Oulu, University of Oulu, Oulu, Finland; Department of Biomedicine, University of Bergen, Bergen, Norway; Department of Chemistry and Interdisciplinary Nanoscience Center (iNANO), Aarhus University, Aarhus, Denmark; Institut Laue-Langevin (ILL), Grenoble, France; Central European Institute of Technology, Masaryk University, Brno, Czech Republic; National Deuteration Facility, The Australian Nuclear Science and Technology Organisation, Locked Bag 2001, Kirrawee DC, NSW 2232, Australia

**Keywords:** myelin, proteolipid protein, DM20, integral membrane protein, small-angle neutron scattering, small-angle X-ray scattering, cryo-EM, synchrotron radiation circular dichroism, atomic force microscopy

## Abstract

The myelin sheath is an essential, multilayered membrane structure that insulates axons, enabling the rapid transmission of nerve impulses. The tetraspan myelin proteolipid protein (PLP) is the most abundant protein of compact myelin in the central nervous system (CNS). The integral membrane protein PLP adheres myelin membranes together and enhances the compaction of myelin, having a fundamental role in myelin stability and axonal support. PLP is linked to severe CNS neuropathies, including inherited Pelizaeus-Merzbacher disease and spastic paraplegia type 2, as well as multiple sclerosis. Nevertheless, the structure, lipid interaction properties, and membrane organization mechanisms of PLP have remained unidentified. We expressed, purified, and structurally characterized human PLP and its shorter isoform DM20. Synchrotron radiation circular dichroism spectroscopy and small-angle X-ray and neutron scattering revealed a dimeric, α-helical conformation for both PLP and DM20 in detergent complexes, and pinpoint structural variations between the isoforms and their influence on protein function. In phosphatidylcholine membranes, reconstituted PLP and DM20 spontaneously induced formation of multilamellar myelin-like membrane assemblies. Cholesterol and sphingomyelin enhanced the membrane organization but were not crucial for membrane stacking. Electron cryomicroscopy, atomic force microscopy, and X-ray diffraction experiments for membrane-embedded PLP/DM20 illustrated effective membrane stacking and ordered organization of membrane assemblies with a repeat distance in line with CNS myelin. Our results shed light on the 3D structure of myelin PLP and DM20, their structure-function differences, as well as fundamental protein-lipid interplay in CNS compact myelin.

## INTRODUCTION

Myelin sheaths are multilamellar membrane wrappings insulating selected neuronal axons, enabling 20-100 times faster conduction of action potentials along myelinated axons. High conduction velocity is fundamental for motor, sensory and cognitive functions. Myelin is formed by specialized glial cells; oligodendrocytes in the central nervous system (CNS) and Schwann cells in the peripheral nervous system (PNS). In the CNS, where larger than 0.2 µm diameter axons are generally myelinated, cell processes of oligodendrocytes spirally wrap around the axons to form an insulating myelin sheath.

Myelin can be divided into two distinct compartments: compact and non-compact myelin. Compact myelin, found in the internodal segments, is a tightly packed multilayered structure, in which the distance between two apposing outer membrane surfaces is only ∼2 nm and no free cytoplasm is present (1). The thickness of the myelin sheath generally depends on the axon diameter; mature CNS myelin may comprise up to 160 membrane turns with a total thickness of 1.7 µm (2). CNS myelin is lipid-rich (70-75% lipid of total dry weight), with a unique lipid profile, mainly consisting of cholesterol (CH), galactosyl ceramide, and ethanolamine plasmalogen (3, 4).

Only a few specific proteins comprise the majority of CNS myelin protein; these include myelin proteolipid protein (PLP), myelin basic protein (MBP), cyclic nucleotide phosphodiesterase (CNPase), myelin oligodendrocyte glycoprotein (MOG) and myelin-associated glycoprotein (MAG). Each of these myelin-specific proteins has a particular function and localization within the myelin sheath. PLP and MBP are the major constituents of CNS myelin, forming 38% and 30% of the total protein mass (5, 6), respectively. Both PLP and MBP exist in the compact compartment and interact tightly with lipid bilayers (7, 8). MBP is a membrane-embedded protein, located on the cytoplasmic leaflet of the myelin membrane, within the major dense line (7, 9). MBP is expressed by both oligodendrocytes and Schwann cells and is crucial for CNS myelin compaction (10). MBP has a major role in myelination, forming a molecular barrier to restrict redundant cytoplasmic and membrane-bound proteins from entering between the myelin lamellae in compact myelin (11).

PLP is a 30-kDa integral membrane protein mainly expressed by CNS oligodendrocytes. Minor expression can be observed in PNS myelin, as well as in kidney distal and proximal tubules (8). PLP is a tetraspan integral membrane protein with cytoplasmic N and C termini. Two alternative isoforms of PLP are expressed; the minor isoform DM20 (26 kDa) lacks 35 residues in its intracellular loop (12). PLP and DM20 are extremely lipophilic and predicted to contain several cysteine-linked fatty acid moieties (13, 14). PLP interacts with membranes and has a high affinity towards CH-rich lipid rafts (15, 16).

PLP is a target for autoantibodies associated with multiple sclerosis (MS) (17–19). Copy-number variation or mutations in the PLP-encoding gene, *PLP1*, result in pathological conditions, such as lethal Pelizaeus-Merzbacher disease (PMD) and milder spastic paraplegia type 2 (SPG2), which both lead to CNS hypomyelination, hypotonia, ataxia, spasticity, and delayed development of motor and cognitive skills (20–23). Additionally, a complete deletion of *PLP1* results in the demyelination of peripheral nerves (24). In transgenic mice, over-expression of *PLP1* produces myelin defects similar to those observed in PMD patients, as well as the accumulation of PLP in vacuoles of the oligodendrocyte soma (25). Respectively, PLP-deficient mice show moderate phenotypic changes with a slightly reduced or delayed myelination of small-diameter axons, increased number of cytosolic channels in compact myelin, inadequate compaction of myelin, and axonal damage (26–29). Hence, a key function of PLP is to adhere myelin lamellae together and enhance the compaction of myelin, thus increasing the physical stability of myelin, as well as to support axon-myelin metabolism.

Due to the high abundance of myelin PLP in brain tissue, PLP has been a target for extensive research for over 70 years (8). Thus far, the challenging extreme hydrophobicity of PLP has hindered its structural and functional characterization. Here, we established a recombinant production system for biologically active human PLP and its shorter isoform DM20. We used small-angle X-ray (SAXS) as well as neutron scattering (SANS), the latter with contrast matching, to reveal the low-resolution 3D structure of dimeric human PLP and DM20 in membrane-mimicking detergent complexes. We demonstrate that lipid membrane-reconstituted recombinant PLP, as well as DM20, induces formation of multilamellar, highly organized membrane assemblies with a repeat distance resembling CNS myelin. The characteristics and determinants of PLP/DM20 membrane stacking were further unraveled using X-ray diffraction, electron microscopy (EM), and atomic force microscopy (AFM). Through our experiments, we provide a novel insight into the structure of two integral membrane proteins of human CNS myelin and the membrane multilayers they assemble together with lipids.

## RESULTS

### Recombinant human PLP and DM20 are dimeric in detergent micelles

Myelin PLP is highly conserved among vertebrates. The human and mouse proteins share 100% identity, and even birds and amphibian proteins are more than 85% identical to mammalian proteins (8). Regardless of the extreme lipophilicity of PLP, we managed to establish a system to produce recombinant human PLP and DM20 in large scale for structural and functional studies. We overexpressed human PLP and DM20 with a cleavable green fluorescent protein (GFP) and an octahistidine tag in the baculovirus-insect cell expression system. PLP and DM20 were solubilized from expression host cell membranes using maltose-based detergents and purified with standard purification techniques. Briefly, after solubilization, immobilized metal affinity chromatography (IMAC) followed by tag cleavage by Tobacco etch virus protease and reverse IMAC were used prior to a final size-exclusion chromatography (SEC) step to obtain pure protein. In SEC, PLP and DM20 (Fig 1A) showed similar elution patterns with one major peak and minor peaks with higher-order oligomers. Pure elution fractions (Fig 1B) were combined from the main peak for further experiments.

**Fig 1.**
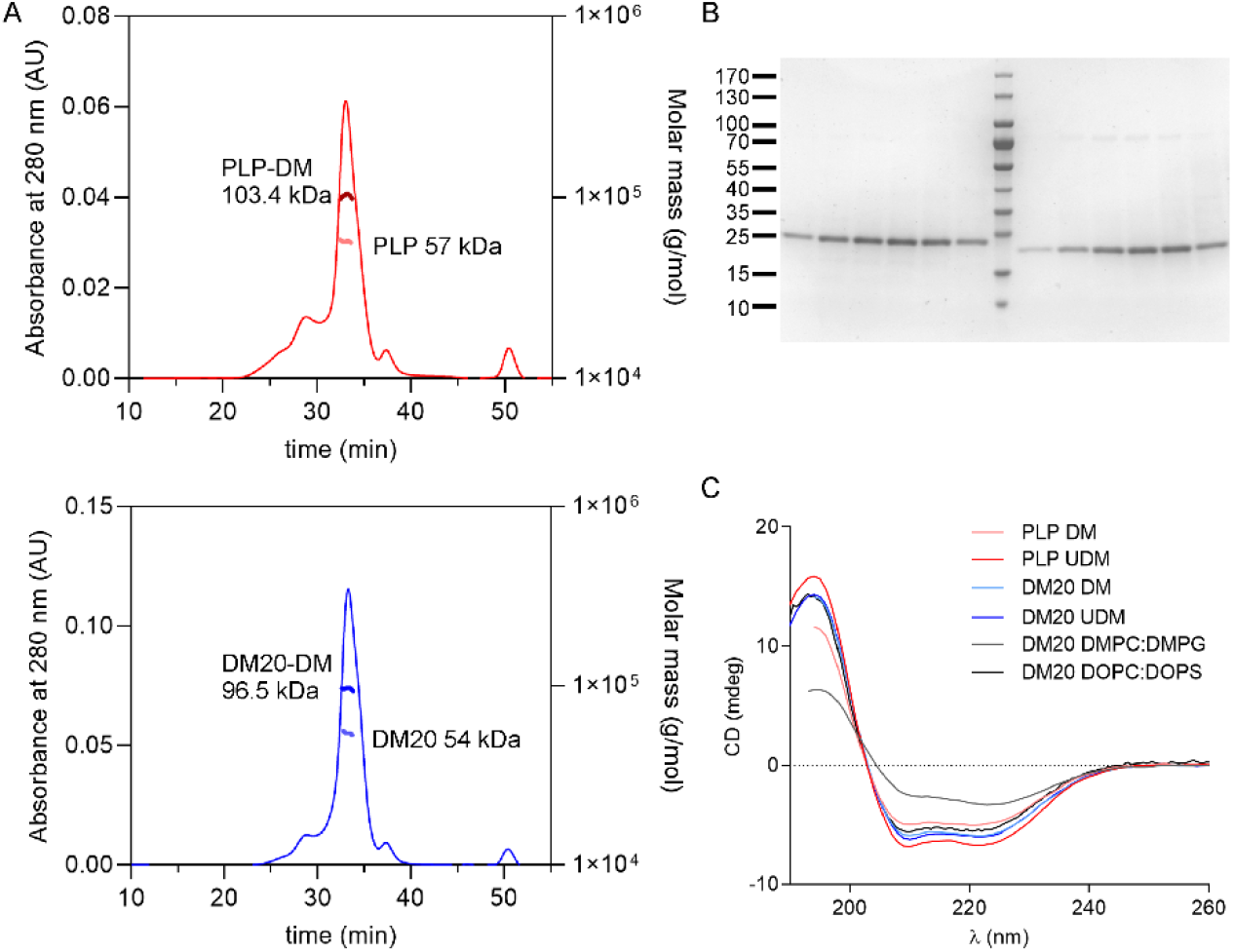
Purification of folded PLP and DM20. A SEC-MALS shows dimeric state for PLP (upper panel) and DM20 (lower panel) in a detergent complex. B The pure PLP and DM20 SEC fractions on Coomassie stained SDS-PAGE. Molecular weight markers in kDa are shown on the left. C SRCD indicates α-helical conformation for PLP and DM20. The lipid composition and detergent tail length slightly affect the protein secondary structure content.

Since PLP and DM20 are transmembrane proteins, they were purified in a complex with a detergent that mimics the membrane bilayer and embeds the hydrophobic transmembrane regions of the protein. This complicates the analysis of the protein oligomeric state, however. To study the oligomeric status of PLP and DM20, the protein molecular weight in detergent complexes, and the amount of bound detergent, were analyzed using multi-angle light scattering (MALS). The total particle molecular weights of the main elution peaks of PLP and DM20, comprising both the protein and bound *n-*decyl*-*β-D*-*maltopyranoside (DM) molecules, were 103.4 kDa and 96.5 kDa, respectively (Fig 1A). The molecular weight of the protein fraction was further determined using protein conjugate analysis, showing a molecular weight of 57.0 kDa for PLP (Fig 1A, upper panel) and 54.1 kDa for DM20 (Fig 1A, lower panel), which is 35 residues shorter. These values are close to the calculated dimeric molecular weights of PLP (60.0 kDa) and DM20 (52.4 kDa), indicating that each protein-detergent complex is composed of two protein molecules surrounded by a detergent belt. Both protein-detergent complexes include ∼45% of detergent, corresponding to 96 and 87 DM molecules for PLP and DM20, respectively.

To follow the folding of human PLP and DM20, we explored the secondary structure content of recombinant PLP and DM20 in complex with maltose-based detergents with either a 10- (DM) or an 11-carbon tail (*n-*undecyl-β-D-maltopyranoside (UDM)) using synchrotron radiation circular dichroism spectroscopy (SRCD). SRCD spectra for both PLP and DM20 indicate a predominantly α-helical conformation, indicating the proper folding of the recombinant proteins (Fig 1C).

A clear difference between detergents was observed for PLP; UDM gave rise to more intense spectral features despite the equal protein concentration in the samples, which can be a sign of more rigid ordering of the transmembrane helices. A minor variation in the spectral shape was detected at 213-222 nm, also possibly reflective of a slightly lower helical content of PLP in DM. On the other hand, DM20 showed no intensity difference between detergents, but spectral shape was slightly altered (at 207-215 nm) when comparing DM20 spectra in DM and UDM; this could relate to minor differences in either helical content or helix packing. These results illustrate an alteration in protein conformation and behavior with different detergent tail lengths and between PLP/DM20. These differences suggest dissimilar physicochemical properties for the two PLP isoforms, including their transmembrane regions that are covered by detergent in these experiments and a lipid bilayer in cells.

### Structure of PLP and DM20 dimers in detergent

Based on the amino acid sequences, both PLP and DM20 comprise four transmembrane helices. Nevertheless, no experimental 3D structural information for PLP or DM20 exists. We measured SEC-SAXS data for human PLP and DM20 in complex with DM and UDM to study their structural characteristics (Fig 2A). Using SEC enabled us to separate empty detergent micelles from protein-detergent complexes. The key SAXS parameters (Table 1) radius of gyration (R_g_), maximum distance in the particle (D_max_), and Porod volume (V_p_) derived from the scattering patterns illustrate the contribution of the detergent belt in the X-ray scattering. The SAXS data indicated slightly larger R_g_, D_max_ and V_p_ for PLP than for DM20, as expected, and the size of the UDM complexes, for both PLP and DM20, were remarkably increased compared to those with a shorter tail DM (Fig 2B, Table 1). All complexes were globular and behaved well in SEC-SAXS.

**Table 1.**
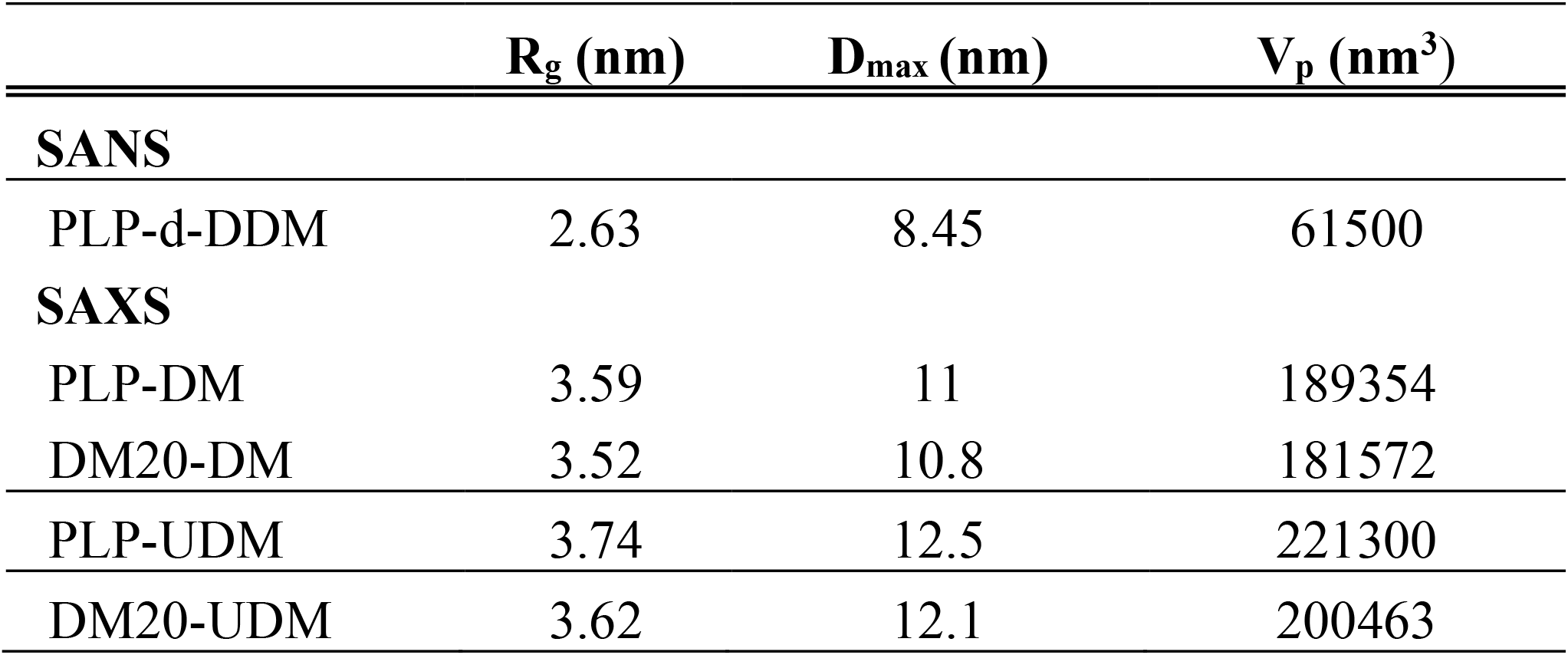
SANS and SAXS parameters.

|  | **R<sub>g</sub> (nm)** | **D<sub>max</sub> (nm)** | **V<sub>p</sub> (nm<sup>3</sup>)** |
| --- | --- | --- | --- |
| **SANS** |  |  |  |
| PLP-d-DDM | 2.63 | 8.45 | 61500 |
| **SAXS** |  |  |  |
| PLP-DM | 3.59 | 11 | 189354 |
| DM20-DM | 3.52 | 10.8 | 181572 |
| PLP-UDM | 3.74 | 12.5 | 221300 |
| DM20-UDM | 3.62 | 12.1 | 200463 |

**Fig 2.**
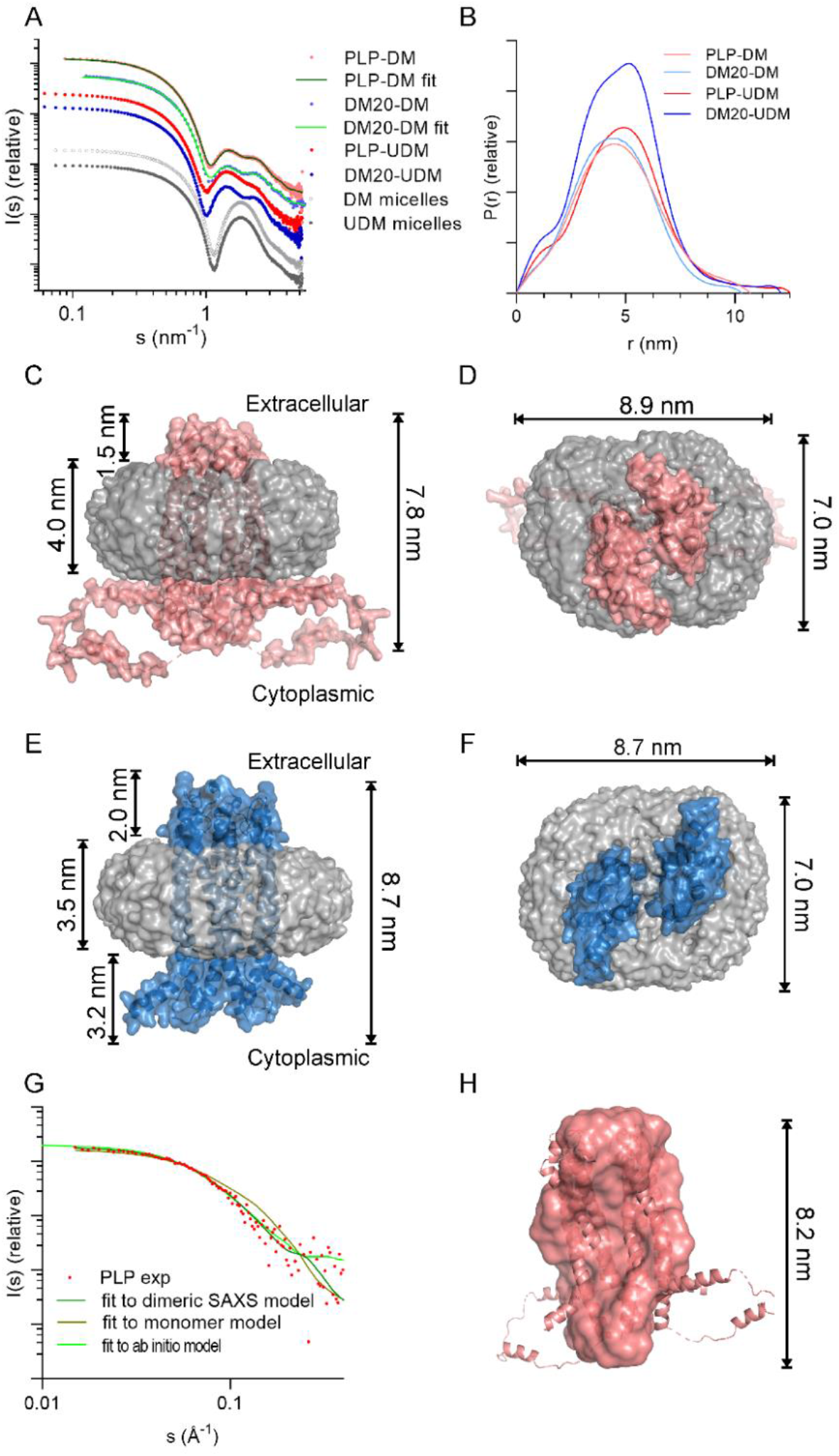
PLP and DM20 are dimeric in detergent. A SAXS patterns of PLP-DM, PLP-UDM, DM20-DM and DM20-UDM complexes as well as empty DM and UDM micelles. The fits of the PLP-DM and DM20-DM models are presented by green lines. B Distance distribution for PLP and DM20 demonstrates the difference in the complex size. C Side view of SAXS model of PLP-DM show 4 nm thick detergent belt around dimeric PLP. The rather long cytoplasmic loop is partially in contact with detergent head groups. The extracellular loops in the intraperiod line of myelin occupy only 1.5 nm space outside the detergent corona. D Top view of PLP-DM model. E Side view of SAXS model of DM20-DM illustrates the shorter cytoplasmic loops and lack of the two helices found in PLP. F Top view of DM20-DM complex. G SANS pattern for PLP-d-DDM complex in D_2_O buffer and fits of the PLP SAXS model and *ab initio* envelope to the experimental SANS data. H The SANS-based model of PLP, superimposed on the SAXS model (cartoons).

To obtain models for protein-detergent complexes based on SAXS data, we utilized monomeric models generated by AlphaFold2 (30, 31) as a starting point in rigid-body modelling. PLP was divided into eight and DM20 into five rigid bodies, and the detergent belt around the protein was modelled with a shell (headgroups) and a core (tails) was modelled around the protein. We were not able to build a model fitting to the scattering pattern with a single copy of PLP or DM20, but two parallel PLP and DM20 molecules surrounded by the detergent corona gave good fits (χ^2^ 4.7 and 3.7, respectively) (Fig 2A). In the PLP-DM complex model (Fig 2C-D), there are 126 DM molecules around the PLP dimer, forming a ∼4 nm thick belt. The detergent belt of the tails covers the hydrophobic surface of PLP, while the intracellular α helices, α1 and α5, are in close proximity to the detergent head groups. The first α helix of the intracellular loop, α4, is not located near the detergent corona and might be more flexible. The extracellular loops of PLP occupy ∼1.5 nm space outside the detergent corona. DM20 lacks most of the intracellular loop (Fig 2E-F), and the detergent belt of the SAXS model of DM20-DM complex was slightly thinner (∼3.5 nm) compared to PLP, although the number of detergent molecules (124) remained nearly the same.

Recently, a novel method to study the structural features of integral membrane proteins in solution was established (32, 33). This includes SEC-SANS in combination with contrast variation and partially deuterated detergent with an aim to render the detergent corona invisible in neutron scattering. We exchanged PLP into a buffer containing chemically modified detergent, with no significant contribution to the scattering pattern in 100% deuterium oxide. The R_g_ and D_max_ (2.65 nm and 8.45 nm, respectively) for PLP from SANS data were remarkably smaller than those measured for PLP-detergent complexes using SAXS (Table 1). This confirms the successful contrast matching of the detergent. The dimeric SAXS model for PLP is in good agreement (χ^2^ = 0.71) with the experimental SANS data (Fig 2G) and further validates our dimeric model, whereas a monomeric model fits more poorly to the SANS data (χ^2^ = 3.00). Additionally, *ab initio* envelopes derived from the SANS data resemble the size and shape of the dimeric SAXS model (Fig 2H) apart from parts of the possibly flexible intracellular regions of PLP.

### PLP and DM20 induce formation of tight multilamellar membrane assemblies

PLP and DM20 are located on lipid bilayers in the compact myelin compartment of CNS where they stabilize multilayered membrane structure (26). CNS myelin is rich in CH and sphingolipids, two major constituents of lipid rafts on cell membranes. We reconstituted PLP and DM20 into unilamellar lipid vesicles with different lipid compositions to explore their function in a more native environment. Notably, most lipid vesicle solutions turned turbid and milky in the presence of PLP or DM20, as previously observed with other myelin proteins (9, 34–36). To investigate this more comprehensively, we recorded the optical density of the samples. Both PLP and DM20, indeed, increased the turbidity of phosphocholine (PC) (18:1) vesicle solutions (Fig 3A). This phenomenon was more pronounced in the presence sphingomyelin (SM) and the most substantial when CH was included in the lipid vesicles. In this experiment, carried out with PC (18:1) vesicles, DM20 had larger effect on the solution turbidity compared to PLP.

**Fig 3.**
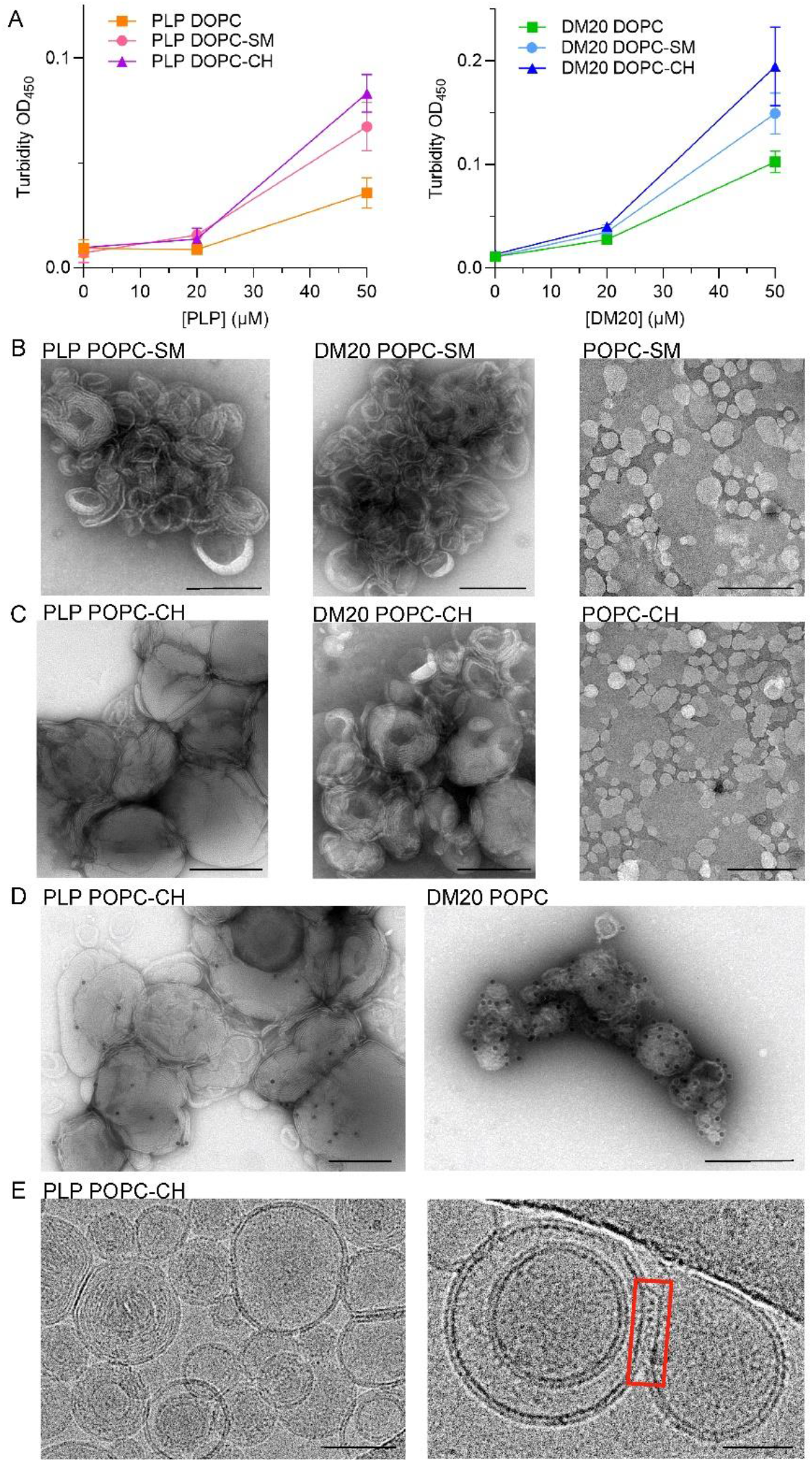
PLP and DM20 induced formation of multilayered membrane assemblies. A PLP (left) and DM20 (right) induce aggregation of PC vesicles. CH and SM enhance the phenomenon, CH having the larger effect. Error bars represent SEM. B Negative stained TEM images of PLP (left) and DM20 (middle) reconstituted PC-SM (B) and PC-CH (C) lipid vesicles and the same vesicles in the absence of protein (right). D Immuno-labelled PLP and DM20 within the membrane aggregates. Scalebars: 200 nm (B-D). E Cryo-EM micrographs of PLP reconstituted in PC-CH vesicles. Red box highlights the visible PLP adhering lipid vesicles together, forming tight junctions between apposing membranes. Scalebar: 50 nm (left panel) 25 nm (right panel).

The conformation of lipid-reconstituted DM20 was investigated by SRCD using PC:phosphatidylserine (PS) (18:1) vesicles, as well as PC:phosphoglyseride (PG) (14:0) vesicles. The SRCD spectrum of DM20 in 18-carbon tail lipid membranes resembles the DM20-detergent spectra, whereas with 14-carbon tail lipid vesicles, DM20 gave a spectrum with lower intensity and clearly altered spectral shape (Fig 1C). This indicates reduced α-helical content or helix packing with shorter lipid tails and agrees with the data obtained from detergent samples.

To study how PLP and DM20 behave when reconstituted in lipid vesicles, and how the morphology of unilamellar lipid vesicles is modified by PLP and DM20, we used EM to image negative stained and flash-frozen samples. In the presence of PLP and DM20, large multilayered membrane aggregates were detected (Fig 3B-E). The dimensions of the membrane aggregates were increased in the presence of CH or SM in PC (16:0-18:1) vesicles (Fig 3B-C). The CH-containing membranes formed regular, tightly packed membrane stacks resembling myelin organization (Fig 3C), while in SM-containing membrane aggregates, the assembly was more irregular and membrane lamellae seemed to be loosely adhered together (Fig 3B). In the absence of PLP and DM20, neither membrane aggregates nor multilayered assemblies were observed (Fig 3B-C, right panels), showing protein-induced formation of such structures. An anti-PLP antibody was utilized in immuno-EM to confirm the presence and successful reconstitution of PLP and DM20 in the lipid aggregates (Fig 3D). The tight packing of CH-containing membrane stacks somewhat hindered the labelling of PLP, while DM20 was more efficiently labelled by the antibody within PC membrane aggregates (Fig 3D).

We further used cryo-EM to study the organization of PLP in PC-CH membrane stacks in more detail. In cryo-EM micrographs, we again observed multilayered lipid bilayer structures glued together by electron dense particles (Fig 3E). The lipid bilayers were often irregular but seemed to contain extra density. Occasionally, PLP could be seen fusing two adjacent bilayers/vesicles together (Fig 3E, right panel). The formed structures had a unique morphology and apparently contained very tight protein-mediated adhesions; these junctions are likely to represent PLP-mediated membrane stacks, resembling those observed in CNS compact myelin (37).

### X-ray diffraction and atomic force microscopy reveal the internal order in membrane stacks

Small-angle X-ray diffraction (SAXD) is a robust technique to study lipid bilayers as well as highly ordered structures (34, 35, 37–39). Ordered systems give rise to Bragg peaks, whose positions in momentum transfer (*s*) can be used to determine repeat distances in the system. We used SAXD to examine PLP/DM20 membrane assemblies, visualized by EM above, to clarify the effect of the protein to lipid (p/l) ratio, lipid composition, as well as putative differences between the isoforms on membrane stacking and repeat distance (Table 2). We observed Bragg peaks at *s* = 0.877-0.939 nm^-1^ corresponding to repeat distances of 6.7-7.3 nm, when PLP or DM20 was reconstituted in PC (16:0-18:1) vesicles. Generally, p/l ratios of 1:500-1:100 produced Bragg peaks; the highest peaks were detected at 1:500 p/l ratio (Fig 4A-H). The samples with higher p/l ratio than 1:100 did not produce any peaks and in the absence of protein, only PC:SM vesicles gave a prominent peak corresponding to a mean repeat distance of 3.7 nm.

**Table 2.** Bragg peak position in SAXD curves and calculated mean repeat distances.

| Sample | 1:500 |  | 1:200 |  | 1:100 |  | Only lipids |  |
| --- | --- | --- | --- | --- | --- | --- | --- | --- |
| **PLP** | 1st peak | 2nd peak | 1st peak | 2nd peak | 1st peak | 2nd peak | 1st peak | 2nd peak |
| POPC | 0.932 (6.74 nm) | no | 0.917 (6.85 nm) | no | 0.877 (7.17 nm) | no | broad | no |
| POPC-SM (4:1) | 0.939 (6.69 nm) | 1.858 | 0.939 (6.69 nm) | 1.892 | 0.877 (7.17 nm) | no | 1.708 (3.67 nm) | no |
| POPC-CH (4:1) | 0.939 (6.69 nm) | 1.863 | 0.939 (6.69 nm) | 1.861 | 0.922 (6.81 nm) | 1.861 | broad | no |
| POPC-SM-CH (8:1:1) | no | no | 0.917 (6.85 nm) | no | 0.913 (6.89 nm) | no | broad | no |
| **DM20** |  |  |  |  |  |  |  |  |
| POPC | 0.932 (6.74 nm) | no | 0.862 (7.29 nm) | 1.709 | 0.862 (7.29 nm) | no | broad | no |
| POPC-SM (4:1) | 0.8783 (7.15 nm) | 1.749 | 0.868 (7.24 nm) | 1.728 | no | no | 1.708 (3.67 nm) | no |
| POPC-CH (4:1) | 0.939 (6.69 nm) | no | 0.939 (6.69 nm) | 1.845 | 0.939 (6.69 nm) | 1.779 | broad | no |
| POPC-SM-CH (8:1:1) | 0.9184 (6.84 nm) | 1.847 | no | no | no | no | broad | no |

**Fig 4.**
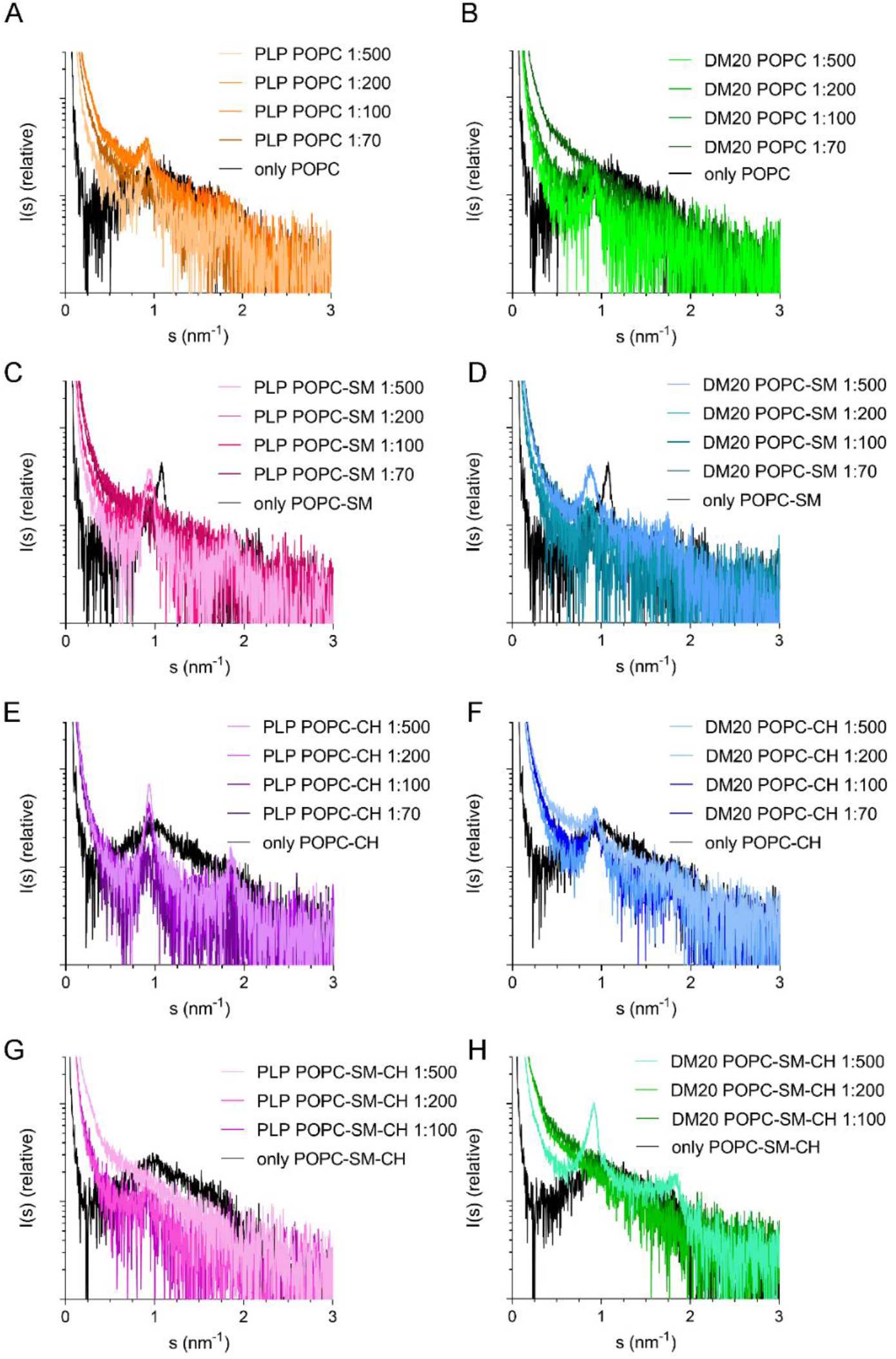
The lipid composition affects membrane stacking by PLP and DM20. A-F SAXD patterns of PLP (A, C, E, G) and DM20 (B, D, F, H) membrane aggregates show Bragg peaks produced by ordered repetitive structures.

With PLP, both SM and CH increased the diffraction peak intensity, CH having a stronger effect. PC-CH lipids with PLP produced clear sharp peaks corresponding to a repeat distance of 6.7-6.9 nm, and with higher PLP density, the mean repeat distance slightly increased. Therefore, the exact spacing between membranes depends on the protein concentration. In PC-SM samples, PLP induced peaks with more variable shape, suggesting a more disordered assembly. However, the position of the Bragg peak, *i*.*e*. the mean repeat distance, remained unaltered.

On the other hand, the mean repeat distance of the DM20:PC-SM membrane aggregates changed in a p/l ratio-dependent manner, whereas that of PC-CH aggregates remained the same. In PC-SM and PC-CH samples, second-order peaks were observed, indicating higher ordering. In the presence of both SM and CH in the PC vesicles, DM20 produced clear sharp peaks with the second order peak with p/l ratio of 1:500, while only minor peaks were detected with PLP in 1:100 and 1:200 p/l ratio. Overall, PLP produces more pronounced Bragg peaks, more ordered and slightly more tightly packed membrane assemblies compared to DM20. This is likely related to the large intracellular loop of PLP affecting membrane adhesion. CH remarkably increased the level of order in PLP membrane assemblies, but interestingly, SM seemed to affect more DM20 than PLP assemblies.

Earlier, we exploited AFM to explore the membrane stacking properties of myelin peripheral membrane proteins (9, 34, 40). Here, we utilized AFM to characterize PLP and DM20 membrane assemblies using PC-SM membranes. Both PLP and DM20 formed multilamellar PC-SM membrane stacks, not observed with PC-SM lipids alone (Fig 5). The formation of multilayered membrane assemblies with a height of >7 nm was slightly more prominent with PLP (Fig 5B-C). Additionally, the thickness of PLP assemblies was generally increased compared to DM20. Clearly, both PLP and DM20 effectively modulated PC-SM membranes and induced the formation of myelin-mimicking multilayered membrane assemblies.

**Fig 5.**
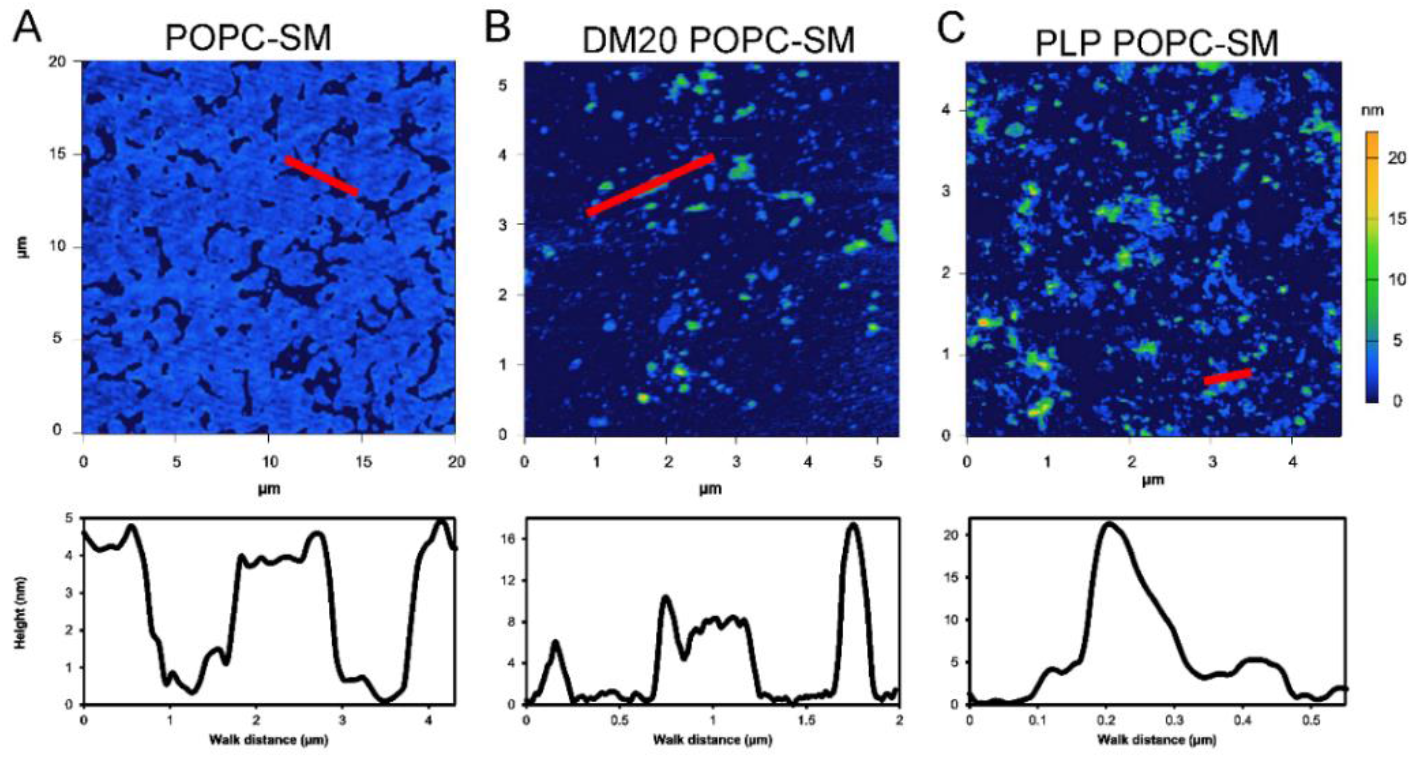
AFM shows the multilayered structure of PLP and DM20 deposited membrane aggregates. A The thickness of a single PC-SM membrane is ∼3.5 nm on mica. B DM20 membrane aggregates have a thickness of 8 nm and above, while PLP produces even thicker structures (C). Red lines represent the walk sections in the images, and the corresponding topology plots are shown in the lower panels.

## DISCUSSION

### Oligomerization of PLP and DM20

CNS compact myelin contains two main protein constituents, PLP and MBP. PLP adheres extracellular membrane leaflets together, but as a transmembrane protein, it may additionally participate in intracellular functions, when the highly ordered multilayered structure is formed (41). PLP is essential to the correct function of the CNS, and *PLP1* defects or alteration in expression patterns lead to severe pathological conditions, such as PMD and SP2 (20, 21, 42). Nevertheless, structural and functional characterization of PLP has remained incomplete.

Using MALS in combination with SAXS and SANS, we determined the low-resolution structure and oligomeric state of PLP and DM20 in membrane-mimicking detergent. Data agrees with both isoforms forming parallel *cis* dimers surrounded by a detergent corona despite the presence of a reducing agent. PLP forms homo-oligomers (dimers and trimers) in oligodendrocyte primary cultures and rat brain lysates, as well as when overexpressed in COS-7 cells, while DM20 mainly exists as a monomer (43, 44). Other tetraspan transmembrane proteins present in compact myelin with a high abundance, such as peripheral myelin protein of 22-kDa (PMP22) and claudins, exist as *cis* dimers on a membrane (45, 46). This suggests that *cis* dimerization is a common characteristic for compact myelin tetraspan integral proteins and putatively involved in the tetraspan protein membrane trafficking, as well as in the organization of multilayered assembly of myelin.

In compact myelin, the adhesion of the extracellular membrane leaflets at the intraperiod line is expected to take place *via* interaction of PLP/DM20 extracellular loops sitting on apposing membranes (47) mediating *trans* oligomerization. The extracellular surface of PLP/DM20 is formed by both positively and negatively charged residues (Fig 6A) and suitable for *trans* oligomerization. The *trans* oligomerization is observed with other compact myelin proteins (48–53), such as myelin P0 and claudins, and seems to be an intrinsic property of these proteins. Another possibility for interactions at the intraperiod line is direct interaction of the PLP extracellular loops with the apposing membrane.

**Fig 6.**
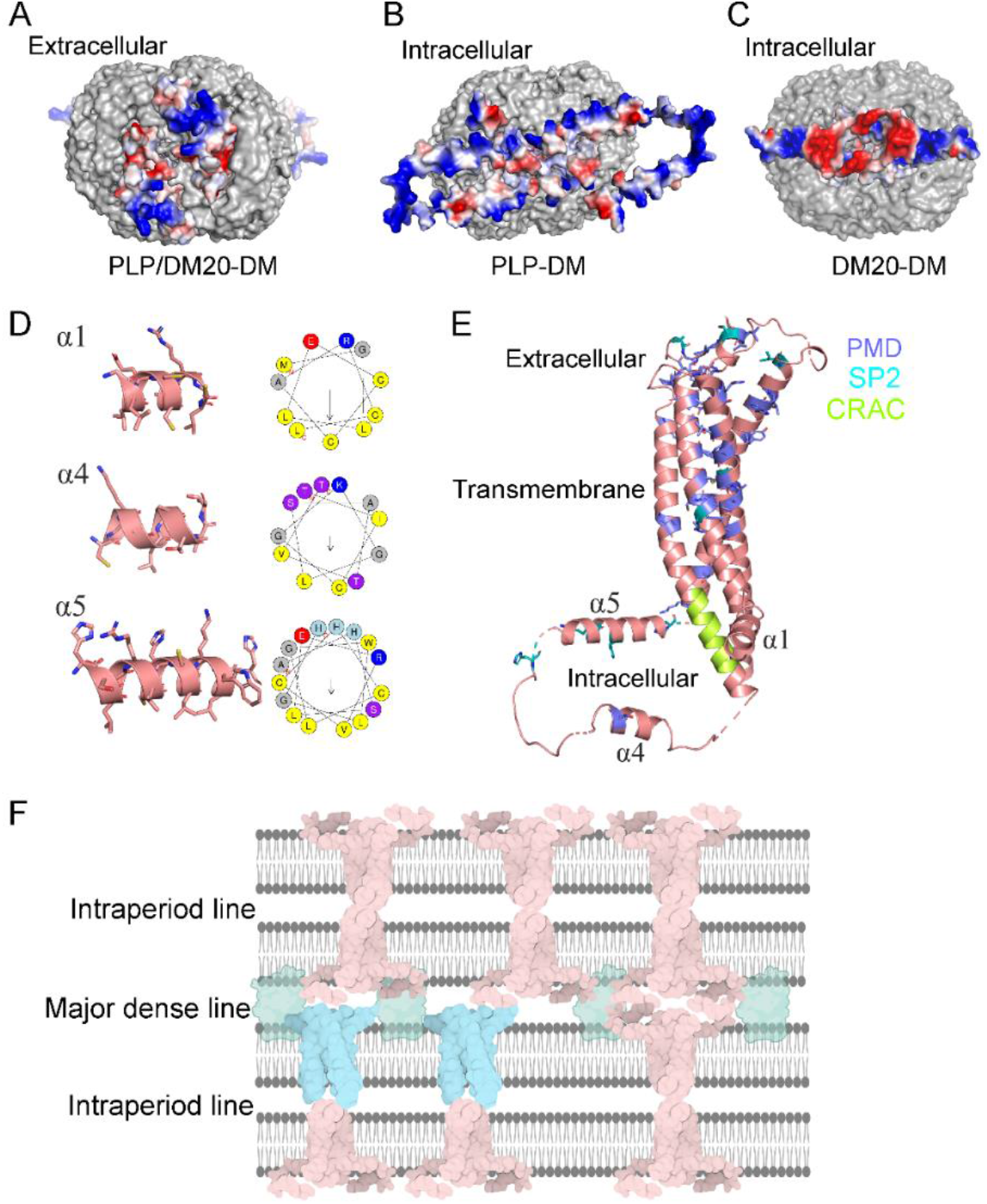
Structural insights. A The vacuum electrostatic surface of PLP-DM from extracellular point of view. B PLP-DM and C DM20-DM models from intracellular point of view illustrate the difference in charge between the two isoforms. D Intracellular α helices of PLP and wheel presentation indicating the amphipathic nature of helix α1 and α5. E The PLP model showing the residues mutated in PMD (purple) or SP2 (cyan), as well as CRAC motif (green). F Schematic view of CNS compact myelin arrangement. The junctions at the intraperiod line represents those seen in the cryo-EM images. PLP (light pink) and DM20 (cyan) presumably form *cis* dimers on myelin membranes. MBP (green) forms the major dense line between the cytoplasmic leaflets. Created with BioRender.com.

The lipids and their modifications on the myelin membrane participate in membrane compaction and myelin stability, by creating an atypical membrane surface with minor repulsive forces (54). This may be a reason for the fact that in our detergent systems, we could not detect significant amounts of higher, so called *trans* oligomers, in which either extracellular loops or intracellular regions of PLP or DM20 would facilitate oligomerization. On the other hand, there are indications that in myelin sheaths, attractive forces gluing the extracellular leaflets together are considerably weaker compared to those on the cytoplasmic membrane leaflet, where MBP adheres the membranes together (41). The ability of PLP/DM20 to stabilize the intraperiod line in compact myelin may arise from a high density of PLP in compact myelin membranes. Weaker attractive forces on the extracellular membrane leaflets may be required during myelin formation, when the membrane layers are wrapped around the axon, and the external membrane leaflets need to be able to move along each other to form multiple myelin layers (41).

### Intracellular regions are membrane-associated

DM20 lacks 35 residues in the intracellular loop of PLP (12), and helices α4 and α5 are absent. This difference alters the surface charge of the intracellular region of PLP (Fig 6B) and DM20 (Fig 6C), DM20 carrying a negatively charged patch. Since cellular membranes generally carry negative surface charge, this difference may affect protein-membrane interactions in the crowded environment of the major dense line. Indeed, mice expressing only the DM20 isoform suffer from reduced myelin stability and disintegration of the myelin membrane layers. These defects became more prominent upon aging (55).

In the PLP-DM SAXS model, helices α1 and α5 are positioned near the detergent head groups. while the third intracellular helix α4 is more distant from the detergent corona. Both α1 and α5 helices have amphipathic character, with one side comprising mainly hydrophobic residues and another side being polar, with positively charged residues (Fig 6D). A PLP-derived peptide containing the second half of the intracellular loop (residues 126-155), *i*.*e*. helix α5, shows a high tendency to fold into an amphipathic helix in the presence of lipid membranes (56). Intriguingly, an overlapping PLP peptide consisting of residues 139-151, with similar behavior (56), has been implicated as a T cell epitope in MS (17) and can be used to generate experimental autoimmune encephalomyelitis, a commonly used mouse model of MS (57). Both helices α1 and α5 located near the detergent corona in our SAXS model contain cysteine residues, which can be acylated with palmitic, oleic, stearic, and palmitoleic acid (13, 58). In both PLP and DM20, in helix α1, all three cysteines, Cys5, Cys6, and Cys9, bind to a fatty acid moiety, whereas in PLP, both cysteines, Cys138 and Cys140, in helix α5 are acylated (58). The acylation pattern of the cysteines validates our conclusion that these helices (α1 and α5) interact with the same membrane layer that the transmembrane helices pass through.

PLP helix α4 is not clearly amphipathic, but it contains a few hydrophobic residues on one side of the helix (Fig 6D). Cys108, located in helix α4, is linked to disulphide-mediated dimerization of PLP, and it is present in both acylated and non-acylated forms *in vivo* (14, 43, 58). This suggests that the first part of the intracellular region, including helix α4, could interact with the apposing membrane layer, and a fatty acid moiety attached to Cys108 could be embedded into this more distal membrane. Unmodified Cys108 might also form a disulfide bond with a PLP counterpart sitting on the apposing membrane. Due to the shorter intracellular loop of DM20, Cys108 or the linked fatty acid may not reach or be able interact with PLP/DM20 or lipids of the apposing membrane layer. This could explain the enhanced ability of PLP over DM20 to stabilize compact myelin structure, as seen in mice expressing only the DM20 isoform (55), as well as the increased cytoplasmic inclusion structures in DM20-overexpressing mice (59). No difference in CNS myelin periodicity was observed in these mice (55). In evolution, PLP appeared in amphibia, reptiles, and birds, when efficient neuronal support and nerve impulse conduction became fundamental for CNS function. PLP could, therefore, play a role in stabilizing the thick myelin sheaths of large vertebrates. DM20 isoform is found in cartilaginous fish, in which the CNS myelin protein composition, in fact, largely resembles vertebrate PNS myelin and no specialization between CNS and PNS myelin proteins has taken place (60, 61).

### Disease-associated mutations in PLP/DM20

Several dysmyelinating conditions are linked to PLP. The duplication as well as more than 60 mutations found in *PLP1* gene lead to PMD, a lethal leukodystrophy (21, 23). PMD is caused by the cytotoxic effect, whereby accumulation of misfolded PLP in ER and oligodendrocytic vacuoles slowly induces oligodendrocyte death that finally leads to axonal swelling and destruction (21). The patient mutations resulting in PMD are clustered in two regions of PLP, in the hydrophobic transmembrane region and the extracellular loops (Fig 6E). These mutations may hinder the proper folding of PLP, reduce the stability of the PLP fold, or affect the PLP trafficking to myelin membranes.

SP2 is a milder PLP-associated leukodystrophy, where progressive degeneration of the axons of upper motor neurons is observed (20). This is the result of the diminished axonal support due to loss of or inadequate function of PLP/DM20 within compact myelin. The sites of the SP2 patient mutations are scattered throughout PLP, in both the extracellular and intracellular loops as well as in the transmembrane regions (Fig 6E). Surprisingly, several mutations are found in the PLP-specific intracellular region missing in DM20. These mutations particularly affect the positively charged residues located in helix α5 and are possibly involved in protein-lipid interactions, illustrating the necessity of helix α5 for PLP function.

### Membrane stacking and cholesterol – PLP interplay

Several myelin-specific proteins have an intrinsic property to be able to modify the morphology of lipid vesicles and adhere membranes to larger membrane aggregates or stacks. This kind of behavior has previously been discovered with the peripheral membrane proteins myelin P2 (35, 36) and MBP (9), as well as with PNS transmembrane proteins PMP22 (62) and P0 (34). In our experiments, recombinant PLP and DM20 revealed a high tendency to assemble into ordered multilayered membranes, in which membrane lamellae are tightly adhered together. Using cryo-EM, we were able to visualize PLP-mediated membrane junctions between the lipid bilayers, which to our knowledge has not been done before. These adhesions are presumably formed via *trans* dimerization of PLP molecules sitting on the apposing bilayers and represent the junctions found in the intraperiod line of CNS compact myelin.

The membrane stacking experiments here were carried out using vesicles containing PCs with relatively long (16:0-18:1) hydrocarbon tails, since the reconstitution of PLP was not possible into vesicles composed of shorter-tail PCs (14:0). However, we were able to reconstitute DM20 also into vesicles with shorter tail lipids (Fig 1D). This detail supports the earlier observation that DM20 is able to adjust its transmembrane regions more flexibly due to the lack of the intracellular hydrophilic region present only in PLP (63).

The ability of PLP to adhere membranes is not charge-driven but more general in nature; in the experiments done using white matter-extracted protein, PLP had similar effects on PC and PC:PS vesicles (64). We observed a clear difference in membrane adhesion when CH or SM was present in the vesicles. CH enhanced the membrane stacking by both PLP and DM20 and induced assembly of larger and more ordered membrane stacks. Both PLP and DM20 contain a CH recognition motif (CRAC) at the end of the second transmembrane helix (Fig 6E). In cellular crosslinking assays, PLP crosslinked with CH but not with other lipids (65). CH and SM together form lipid rafts on the plasma membrane, and these microdomains are highly enriched in myelin. PLP has a high affinity towards these membrane microdomains and especially towards CH, which is required for PLP raft association (16, 65). Interestingly, CH is fundamental in the early stages of axon wrapping, and blocking CH synthesis halts myelin wrapping and compaction (66, 67). None of the *PLP1* disease mutations reported is located in the CRAC motif (Fig 6E). Our findings illustrate the significance of the interplay between membrane lipids and myelin proteins in the complex process of myelination.

### Mean repeat distances of myelin-like assemblies

During decades of intensive research, X-ray diffraction experiments on myelinated nerves have revealed typical diffraction patterns as well as intraperiod distances for PNS and CNS myelin (37–39, 68–71). Here, with reconstituted recombinant human PLP, the measured repeat distance in a multilayer was 6.7-7.2 nm, depending on the p/l ratio and the presence of SM and CH in the lipid membrane. The high abundance of PLP on the membrane, *i*.*e*. a high p/l ratio, occupies more space between the membranes and increases the mean repeat distance. The thickness of a PC (16:0-18:1) bilayer is 4.2 nm, while CH and SM slightly increase the thickness (72). This indicates that the intermembrane space occupied by PLP varies between 2.2-3.0 nm. It must be noted that the orientation of PLP/DM20 molecules on the reconstituted membranes is presumably random, and we cannot differentiate the occupied space of extra and intracellular domains. Tissue-extracted myelin P0, a protein responsible for extracellular adhesion of PNS myelin membranes, forms zipper-like structures with a 5-nm intermembrane space at the intraperiod line, when reconstituted into PC membranes (34). This difference between PLP and P0 is in good agreement with the SAXD measurements carried out using myelinated nerves derived from nervous systems. In the PNS, the variation between specimens is smaller and the intraperiod distance is 18.4 nm. In CNS myelin, the intraperiod distance varies between 16.0-16.5 nm (73, 74). Based on cryo-EM micrographs, the PNS-specific tetraspan protein PMP22 shows tighter packing of membrane aggregates compared to P0 but correspondingly less compact compared to PLP or DM20 (62).

In the CNS, the intraperiod line is adhered by PLP, while the intracellular major dense line is formed and irreversibly formed by MBP. MBP deficiency leads to demyelinating neuropathy affecting mainly CNS (75, 76). Using the peripherally attached cytosolic proteins P2, MBP, and the cytoplasmic domain of P0, we have measured repeat distances in the 7-8 nm range (9, 34, 36). These experiments revealed a PNS major dense line-like structure with a distance of 3-4 nm between apposing membranes. The SAXD experiments with MBP membrane stacks revealed a mean repeat distance of 8 nm, when PC:PG (14:0) were used, meaning that MBP occupies a 3.5-4.0 nm space between the lipid bilayers (9). MBP and PLP would then jointly produce a myelin intraperiod distance exactly matching the one measured in myelinated CNS nerves (16.0-16.6 nm) (Fig 6F).

### Concluding remarks

Our work represents the first structural characterization of human PLP/DM20 and the supramolecular assemblies they form together with lipid bilayers. However, experiments with membrane systems containing all main protein components of CNS compact myelin remain to be accomplished. In the future, studies on such myelin-mimicking model systems are needed to elucidate the complex interplay between lipids and proteins in myelin formation, maintenance and remodeling, as well as in demyelinating disorders. Our work provides tools towards both high-resolution structural analysis of myelin-like membranes as well as the formation of multicomponent systems for a full understanding of the molecular structure and dynamics involved in compact myelin formation and stability.

## MATERIALS AND METHODS

### Expression and purification of PLP and DM20

PLP and DM20 were expressed using the Bac-to-Bac Baculovirus expression system (Thermo Fisher Scientific) in *Spodoptera frugiperda* 21 (*Sf*21) cells. PLP (acc. NM_000533) and DM20 genes were cloned into pFastBac dual vector (Thermo Fisher Scientific) with a cleavable C-terminal eGFP-His_8_ tag. The PLP/DM20 plasmid was transformed into DH10Bac strain (Thermo Fisher Scientific) for bacmid production. *Sf*21 insect cells were transfected with PLP/DM20 bacmid using Fugene 6 transfection reagent (Promega Corp.) and baculoviruses were collected and used for preparation of a high-titer virus stock. *Sf*21 cells were infected using PLP/DM20 virus stock and cells were harvested 72-90 h after the infection and washed with phosphate buffered saline (137 mM NaCl, 2.7 mM KCl, 8 mM Na_2_HPO_4_, and 2 mM KH_2_PO_4_). Cells were suspended in lysis buffer containing 0.5 M NaCl, 0.5 mM TCEP, 10% glycerol, 50 mM Tris pH 7.5 and manually broken using dounce homogenizer in the presence of 20 µg/ml DNase I (AppliChem) and SigmaFast protease inhibitor cocktail tablets (Sigma-Aldrich). The membrane fraction was separated with ultracentrifugation (1 h, +4 °C, 234788 g). PLP and DM20 were solubilized using 1% (w/v) UDM or 1.25% (w/v) DM in lysis buffer with a gentle agitation at +4 °C for 2 h. Insoluble material was removed by a second ultracentrifugation step (1 h, +4 °C, 234788 g). The supernatant was mixed and rotated with HisPur NiNTa resin (Thermo Fisher Scientific) for 1 h at +4 °C. After washing with washing buffer (0.06% UDM/0.2% DM, 50 mM imidazole, 0.5 M NaCl, 0.5 mM TCEP, 10% glycerol, 50 mM Tris pH 7.5), PLP/DM20 was eluted with the elution buffer containing 0.06% UDM/0.2% DM, 250 mM imidazole, 0.3 M NaCl, 0.5 mM TCEP, 10% glycerol, 20 mM Tris pH 7.5. The imidazole was removed, and the tags were cleaved by Tobacco Etch virus protease in dialysis against SEC buffer. The sample was passed through the HisPur resin to remove eGFP-His_8_ tag and TEV protease and further purified using Superdex 200 Increase 10/300 gel filtration column (GE Healthcare) in SEC buffer containing 0.06% UDM /0.2% DM, 0.2-0.3 M NaCl, 0.5 mM TCEP, 2-10% glycerol, 20 mM Tris pH 7.5. PLP/DM20 was concentrated using Sartorius Vivaspin 20 concentrator (MWCO 50-100 kDa) to appropriate concentration.

### Protein conjugate analysis by multi-angle light scattering

Molecular masses of PLP and DM20 detergent complexes were determined using SEC-MALS at the core facility for Biophysics, Structural Biology, and Screening (BiSS) at the University of Bergen. A Superdex 200 Increase 10/300 (GE Healthcare) column was used to separate empty detergent micelles from the protein-detergent complexes. 50 µg of PLP or DM20 in SEC buffer (0.2% DM, 0.3 M NaCl, 0.5 mM TCEP, 5% glycerol, 20 mM Tris pH 7.5) was injected into the SEC column using Shimadzu Prominence I LC-2030C HPLC unit, and UV absorption was recorded by LC-2030/2040 PDA detector (Shimadzu). The light scattering was measured by a Wyatt miniDAWN TREOS instrument (Wyatt Technology) and refractive index was determined using a RefractoMax 520 detector (ThermoScientific). Data were analysed using Astra 7 (Wyatt Technology) utilizing protein conjugate analysis with dn/dc values of 0.1473 ml/g and 0.1850 ml/g for DM and proteins, respectively.

### Synchrotron radiation circular dichroism spectroscopy

SRCD spectra between 170 and 280 nm were recorded using a 100‐μm quartz cuvette (Suprasil, Hellma Analytics), on the AU‐CD beamline at the ASTRID2 synchrotron storage ring (ISA, Aarhus, Denmark). 0.5 mg/ml of PLP and DM20 in buffer containing 0.06% UDM or 0.2% DM, 150 mM NaF, 0.25 mM TCEP, 2% glycerol, 20 mM Tris pH 7.5 were used in all measurements. The measurements were carried out at +30 °C.

### Small-angle X-ray scattering

SAXS measurements were performed at beamline SWING, SOLEIL (Paris, France) (77) and at EMBL/DESY beamline P12, PETRA III (Hamburg, Germany) (78). The protein concentrations of 4.2-5.2 mg/ml for PLP and 4.9-6.2 mg/ml for DM20 were used in SEC-SAXS measurements. SEC was carried using a Superdex 200 Increase 10/300 column (GE Healthcare), a 0.1 ml/min flow rate, with a buffer containing 0.06% β-UDM /0.2% DM, 0.3 M NaCl, 0.5 mM TCEP, 3% glycerol, 50 mM Tris pH 7.5. All SAXS measurements were done at +10-20 °C. Data were processed and analyzed using Foxtrot or ATSAS (79). SAXS data for PLP and DM20 are available using doi: 10.5281/zenodo.6324818.

The SAXS data were modelled by methods similar to those described in (80–82), where scattering length densities and molecular properties are used as restraints. The structure of the proteins were predictions by AlphaFold2 (30, 31). The rim of DM covering the hydrophobic belt of the proteins were modelled using Monte Carlo generated points. Initially, points randomly and homogeneously distributed in a box are generated and geometrical restraints are used for selecting points to present the hydrocarbon tails and the maltoside headgroups of DM, which have distinctly different electron densities; that of the hydrocarbon tail is lower than that of the buffer and that of the headgroups is higher. The surfactant rim was generated to decorate the surface of the protein. This was done by placing the protein with the trans-membrane axis along the *z* axis, so that it cuts the *x-*y plane at *z* = 0. The excluded volume of the protein is then calculated from the atomic positions as a set of points on a grid allowing the atoms to occupy a certain number of grid points. Next, the surface grid points of the excluded volume are located and the *z* = 0 points are found. This gives a set of points at the circumference of the protein, which lie on a closed curve, and when the protein is appropriately positioned, the curve can be at center of the hydrophobic region of the protein. Monte Carlo points that have a minimum distance smaller than *R*, to any point on the closed curve, and which are not overlapping with the protein within 3.0 Å, are selected for representing the hydrocarbon tails of the DM. Similarly, headgroup points are selected as points with a minimum distance to the points on the closed curve between *R* and *R* + *D*, where *D* is the width of the headgroup layer; only points not included in the hydrocarbon rim and not overlapping with the protein are used. For ensuring that the headgroup and tails are connected, it is furthermore checked that the minimum distance between a headgroup point and any hydrocarbon point is not larger than *D*. With this approach, the detergent core-shell structure is in principle described by only two parameters. However due to the curvature of the tail-headgroup interface, it was decided to allow the core of the rim and the shell of the rim to have different radius and shell thickness in, respectively, the *x-y* and the *z* directions, thus increasing the number of parameters to four. A large curvature may ‘dilute’ the headgroups and lead to a lower scattering contrast of the shell where the curvature is large, however, an easy way to implement this effect was not found. As thickness of the shell and scattering contrast correlates significantly, it was decided to have a variation of the shell thickness instead.

The tail rim is assumed not to contain any hydrating water molecules and therefore its volume can be calculated from the original density of points in the box and the number of points selected from the full set. The number of DM molecules is obtained by dividing the hydrocarbon volume by the volume of a decyl chain, 294 Å^3^. The electron density of this buffer was estimated to be 0.340 e/Å^3^. The standard scattering length density of the protein of 2.00 × 10^10^ cm/g was modified accordingly and also the electron density of surfactant tails and headgroups were calculated taking this into account. For the ‘dry’ volume a maltoside headgroup a value of 350 Å^3^ was used. In the model, there is no restraint on *D*, (except that it should respect a physically reasonable value so that it is less than 10 Å), and therefore hydrating water molecules are implicitly allowed to be present within the headgroup shell. The implementation respects that the total excess scattering length of the core and the shell, respectively, corresponds to a specific number of DM molecules, as calculated from the core volume. The total scattering length of core and shell were distributed on the Monte Carlo points representing, respectively, the core and the shell. A layer of hydrating dummy atoms was added around the protein, where it is not covered by surfactant, to represent a hydration layer. It was assigned a typical value as used for other proteins (83).

As an initial test, the modelling approach without including a protein was applied to pure micelles of DM from the same SEC-SAXS run. The procedure gave satisfactory fits to the SAXS data, thus confirming that the molecular restraints for DM were correctly implemented.

In agreement with the MALS results, it turned out to be impossible to obtain reasonable fits for only one protein molecule per complex and therefore two molecules were included with P2 symmetry in a parallel configuration. The position of the protein was optimized together with the parameters describing the rim. A steric overlap penalty function was added to the reduced chi-squared with a certain weight, to avoid overlap between the two proteins (83). The optimization was done making random searches initially with larger steps, which gradually were decreased. Both the position of the protein and the parameters describing the rim was optimized. As a Guinier plot revealed an upturn at low *q*, due to some aggregation, data below 0.02 Å^-1^ were omitted for both data sets. During the optimization, the parameter values for the best fit with the lowest value of the sum of chi-squared and the penalty function was kept. The approach gave reasonable fits, however, in order to improve them and take into consideration the uncertainty in the structure prediction in particular for the flexible loops, the structures were divided into 5 bodies for the DM20 construct and 8 bodies for PLP. Additional harmonic restraints for the Cα atoms at the connections, around the ideal values of 3.8 Å, was added to chi-squared and the overlap penalty function to keep the structures connected.

### Small-angle neutron scattering

The SANS measurements were carried out in combination with an *in situ* SEC system at beamline D22 at Institute Laue-Langevin, (Grenoble, France). PLP in D_2_O buffer (0.2% DM, 0.3 M NaCl, 0.25 mM TCEP, 20 mM Tris pH 7.5, ∼100% D_2_O) was slowly passed through Superdex 200 Increase 10/300 column (GE Healthcare) to exchange protonated DM to partially deuterated d-DDM (tail 89 % and head 57% deuterated) using a buffer containing 0.04 % d-DDM (The National Deuteration Facility at the Australian Nuclear Science and Technology Organization (ANSTO)), 0.3 M NaCl, 0.25 mM TCEP, 20 mM Tris, pD 7.4 in ∼100% D_2_O). The SANS data was collected while the sample was eluted from SEC column. A neutron wavelength of 6 Å ± 10 % at sample-detector distances of 2 m and 5.6 m, respectively, were used to cover the s-range from 0.009 to 0.7 Å^−1^. The temperature of the sample cells was set to 10 °C. Data were processed and analysed using GRASP package. The software CRYSON (84) were used to calculate theoretical SANS curves on structural models and fit them to the experimental data. The *ab initio* modelling was carried by GASBOR (85). Original data are available using doi: 10.5291/ILL-DATA.8-03-1006 and 10.5291/ILL-DATA.8-03-1022.

### Lipid reconstitution and turbidity measurements

Unilamellar lipid vesicles containing 10 mM DMPC:DMPG (1:1), 6.35 mM DOPC, DOPC:SM (4:1), DOPC:CH (4:1) or 6.58 mM POPC, POPC:SM (4:1), POPC:CH (4:1) and POPC:SM:CH (8:1:1) were prepared as previously described (9, 56). Briefly, lipids were dissolved in chloroform:methanol (4:1) mixture or in chloroform in using molar ratios. The solvent was evaporated, and membranes were dissolved in water and sonicated until the vesicle solution became clear. An appropriate amount 1-2.5 mM of unilamellar vesicles with 0.4 % DM were mixed with PLP or DM20 purified with DM as a detergent using the protein-to-lipid ratios of 1:35-1:500. The protein-lipid mixtures were incubated for 1 h at +22 °C and then transferred into Slyde-A-Lyzer mini dialysis devices (Thermo Fisher Scientific) with 20 kDa MW cut-off or 10-20 µl dialysis buttons (Hampton Research) covered with SpectraPor dialysis membrane (MW cut-off 12-14 kDa). To remove DM and reconstitute PLP and DM20 into lipid membranes, samples were dialyzed against reconstitution buffer containing 0.2-0.3 M NaCl, 0.5 mM TCEP, 0-3% glycerol, 20-50 mM Tris pH 7.5 at +22 °C with gentle shaking for 5-7 days. The buffer was changed daily. For turbidity measurements, 0-50 µM PLP or DM20 was reconstituted in 0.5 mM lipid vesicles as described before. The turbidity of the solution was measure by recording OD at 450 nm using NanoDrop 1000 Spectrophotometer (ThermoFisher Scientific). The measurements were done as triplicates.

### Negative staining, immunoelectron microscopy and electron cryomicroscopy

For negative staining, protein-lipid stacks obtained using 5 µM PLP/DM20 and 1 mM unilamellar vesicles (p/l ratio 1:200) containing POPC, POPC:SM (4:1) or POPC:CH (4:1) were used. In control samples only lipid vesicles were included but the sample treatment was the same. 4-µl samples were pipetted onto glow-discharged carbon-coated copper grids and incubated for 1 min. The excess solution was removed using filter paper. After 4 washes in dH_2_O droplets, samples were negatively stained with two drops of 2% uranyl acetate for 12 s in each and air-dried. In immuno-EM, anti-myelin PLP antibody (Abcam, 105784) was used as a primary antibody and detected by protein A conjugated gold particles (10 nm). The samples were then negatively stained using 2% uranyl acetate as described above. TEM images were recorded using a Tecnai G2 Spirit 120 kV instrument equipped with a Quamesa CCD camera (Olympus Soft Imaging Solutions) at the EM core facility of Biocenter Oulu.

Cryo-EM experiments were carried out at Cryo-Electron Microscopy and Tomography Core Facility, CEITEC MU/ CIISB (Brno, Czech Republic). For cryo-EM, PLP membrane stacks with p/l ratio of 1:200 and 2 mM POPC-CH unilamellar vesicles were used. Samples were applied to glow-discharged, holey copper grids (QUANTIFOIL R 2/2). 3.5-μl samples were adsorbed for 1 min at +20 °C, 95% humidity. Grids were blotted for 8 s from both sides and vitrified by plunging into liquid nitrogen–cooled liquid ethane using a Vitrobot mark IV (ThermoScientific). The grids were imaged using Talos Arctica transmission electron microscope operated at 200 kV. Micrographs were collected at calibrated pixel size of 1.23 A/px on direct electron detector Falcon 3EC (ThermoScientific) operating in charge integration mode. The exposure time was 1s and overall dose was 40 e/Å^2^.

### Small-angle X-ray diffraction

For SAXD measurements, PLP and DM20 were reconstituted in 1 mM unilamellar vesicles consisting of POPC, POPC:SM, POPC:CH or POPC:SM:CH. The protein-lipid molar ratios of 1:70, 1:100, 1:200 and 1:500 were used. The control samples contained only lipids but were handled similarly. The SAXD experiments were carried out at EMBL beamline P12 (PETRA-III/DESY) (78) and at SWING (SOLEIL) (77) at 20 °C. The Bragg peak positions in momentum transfer *s* were used to calculate the mean repeat distance in the membrane stacks using the formula *d* = 2π/*s*, where *d* is the repeat distance and *s* is defined as *s* = 4πsinθ/λ, where θ is the scattering angle and λ is the X-ray wavelength. Data were processed using ATSAS (79). SAXD data are available using doi: 10.5281/zenodo.6324818.

### Atomic force microscopy

10-15 µM PLP/DM20 and 394-526 µM POPC-SM vesicles were used in lipid reconstitutions for AFM measurements. The controls contained only POPC:SM vesicles. Freshly cleaved ruby muscovite mica (1.2 cm diameter) was used as substrate for depositing lipid and protein-lipid samples. Each sample was diluted to reach a 260-500 µM lipid concentration in 20 mM HEPES, 150 mM NaCl, pH 7.5. To each lipid and protein-lipid sample, 2 mM and 5 mM CaCl_2_ were added, respectively. The mica was covered entirely with 100 – 200 µl of sample for each deposition. Lipid samples were incubated against mica for 20 min at +30 °C and protein-lipid samples were incubated overnight at ambient temperature. During deposition, the mica was covered with a Petri dish lid to minimize evaporation. After this, the sample suspension was removed by suction using paper and washed twice with the same buffer including the appropriate CaCl_2_ concentration. The mica was covered with the same buffer lacking CaCl_2_. AFM imaging was performed immediately after the deposition procedure. All samples were prepared in duplicate.

An Asylum Research MFP-3D Bio instrument was used for AFM imaging at the solid-liquid interface. All images were acquired at ambient temperature. Acquisition control and image processing was performed using Igor Pro 6.37. Olympus OMCL-TR800PSA cantilevers with spring constants between 0.628 – 0.657 N m^-1^ and resonance frequencies between 75.732 – 76.465 kHz were used in alternative current mode at a 90° scan angle. Square scans were acquired from areas between 4.5 – 20 µm. For lipid depositions, 256 × 256 pixel images were acquired at scan speed of 0.8 Hz. For protein-lipid depositions, 512 × 512 pixel images were acquired at a scan speed of 0.6 Hz. At least 3 sections were scanned for each deposition to check for deposition heterogeneity.

## ACKNOWLEDGEMENTS

This work was funded by the Academy of Finland, grant number 275225 and Jane and Aatos Erkko Foundation. Beamtime and user support at EMBL/DESY, SOLEIL, ILL and ISA are gratefully acknowledged. Travel to synchrotrons was supported by the European Union Horizon 2020 programs iNEXT (Grant 653706) and CALIPSOplus (Grant 730872). The National Deuteration Facility at ANSTO is partly funded by The National Collaborative Research Infrastructure Strategy (NCRIS), an Australian Government initiative. We acknowledge the cryo-electron microscopy and tomography core facility CEITEC MU of CIISB, Instruct-CZ Centre supported by MEYS CR (LM2018127). The use of the facilities and expertise of the Biocenter Oulu Protein analysis, Structural Biology and EM core facilities is thankfully acknowledged. We further acknowledge the use of the Core Facility for Biophysics, Structural Biology, and Screening (BiSS) at the University of Bergen.

## REFERENCES

1. R. M. Stassart, W. Möbius, K.-A. Nave, J. M. Edgar, The Axon-Myelin Unit in Development and Degenerative Disease. Front Neurosci 12, 467 (2018).

2. C. Hildebrand, S. Remahl, H. Persson, C. Bjartmar, Myelinated nerve fibres in the CNS. Prog Neurobiol 40, 319–384 (1993).

3. S. Schmitt, L. C. Castelvetri, M. Simons, Metabolism and functions of lipids in myelin. Biochimica et biophysica acta 1851, 999–1005 (2015).

4. K.-A. Nave, H. B. Werner, Myelination of the nervous system: mechanisms and functions. Annu Rev Cell Dev Biol 30, 503–533 (2014).

5. O. Jahn, S. Tenzer, H. B. Werner, Myelin proteomics: molecular anatomy of an insulating sheath. Molecular neurobiology 40, 55–72 (2009).

6. O. Jahn, et al., The CNS Myelin Proteome: Deep Profile and Persistence After Post-mortem Delay. Front. Cell. Neurosci. 14 (2020).

7. J. M. Boggs, Myelin basic protein: a multifunctional protein. Cell Mol Life Sci 63, 1945–1961 (2006).

8. J. M. Greer, M. B. Lees, Myelin proteolipid protein–the first 50 years. The international journal of biochemistry & cell biology 34, 211–215 (2002).

9. A. Raasakka, et al., Membrane Association Landscape of Myelin Basic Protein Portrays Formation of the Myelin Major Dense Line. Scientific reports 7, 4974-017-05364–3 (2017).

10. C. Readhead, L. Hood, The dysmyelinating mouse mutations shiverer (shi) and myelin deficient (shimld). Behav Genet 20, 213–234 (1990).

11. S. Aggarwal, et al., A size barrier limits protein diffusion at the cell surface to generate lipid-rich myelin-membrane sheets. Dev Cell 21, 445–456 (2011).

12. K. A. Nave, C. Lai, F. E. Bloom, R. J. Milner, Splice site selection in the proteolipid protein (PLP) gene transcript and primary structure of the DM-20 protein of central nervous system myelin. Proc Natl Acad Sci U S A 84, 5665–5669 (1987).

13. T. Weimbs, W. Stoffel, Proteolipid protein (PLP) of CNS myelin: positions of free, disulfide-bonded, and fatty acid thioester-linked cysteine residues and implications for the membrane topology of PLP. Biochemistry 31, 12289–12296 (1992).

14. O. A. Bizzozero, L. K. Good, J. E. Evans, Cysteine-108 is an acylation site in myelin proteolipid protein. Biochem Biophys Res Commun 170, 375–382 (1990).

15. H. B. Werner, et al., A critical role for the cholesterol-associated proteolipids PLP and M6B in myelination of the central nervous system. Glia 61, 567–586 (2013).

16. I. Castello-Serrano, J. H. Lorent, R. Ippolito, K. R. Levental, I. Levental, Myelin-Associated MAL and PLP Are Unusual among Multipass Transmembrane Proteins in Preferring Ordered Membrane Domains. J Phys Chem B 124, 5930–5939 (2020).

17. J. M. Greer, M. P. Pender, Myelin proteolipid protein: an effective autoantigen and target of autoimmunity in multiple sclerosis. Journal of Autoimmunity 31, 281–287 (2008).

18. N. C. Cloake, J. Yan, A. Aminian, M. P. Pender, J. M. Greer, PLP1 Mutations in Patients with Multiple Sclerosis: Identification of a New Mutation and Potential Pathogenicity of the Mutations. Journal of clinical medicine 7, 10.3390/jcm7100342 (2018).

19. J. M. Greer, E. Trifilieff, M. P. Pender, Correlation Between Anti-Myelin Proteolipid Protein (PLP) Antibodies and Disease Severity in Multiple Sclerosis Patients With PLP Response-Permissive HLA Types. Front. Immunol. 11 (2020).

20. C. Blackstone, Hereditary spastic paraplegia. Handb Clin Neurol 148, 633–652 (2018).

21. K. Inoue, Pelizaeus-Merzbacher Disease: Molecular and Cellular Pathologies and Associated Phenotypes. Adv Exp Med Biol 1190, 201–216 (2019).

22. K. A. Lüders, J. Patzig, M. Simons, K.-A. Nave, H. B. Werner, Genetic dissection of oligodendroglial and neuronal Plp1 function in a novel mouse model of spastic paraplegia type 2. Glia 65, 1762–1776 (2017).

23. R. Singh, D. Samanta, “Pelizaeus-Merzbacher Disease” in StatPearls, (StatPearls Publishing, 2021) (May 24, 2021).

24. J. Y. Garbern, et al., Proteolipid protein is necessary in peripheral as well as central myelin. Neuron 19, 205–218 (1997).

25. S. A. Karim, et al., PLP overexpression perturbs myelin protein composition and myelination in a mouse model of Pelizaeus-Merzbacher disease. Glia 55, 341–351 (2007).

26. M. Klugmann, et al., Assembly of CNS myelin in the absence of proteolipid protein. Neuron 18, 59–70 (1997).

27. J. Rosenbluth, K.-A. Nave, A. Mierzwa, R. Schiff, Subtle myelin defects in PLP-null mice. Glia 54, 172–182 (2006).

28. I. Griffiths, et al., Axonal swellings and degeneration in mice lacking the major proteolipid of myelin. Science 280, 1610–1613 (1998).

29. H. Jurevics, et al., Normal metabolism but different physical properties of myelin from mice deficient in proteolipid protein. J Neurosci Res 71, 826–834 (2003).

30. J. Jumper, et al., Highly accurate protein structure prediction with AlphaFold. Nature 596, 583–589 (2021).

31. M. Varadi, et al., AlphaFold Protein Structure Database: massively expanding the structural coverage of protein-sequence space with high-accuracy models. Nucleic Acids Research 50, D439–D444 (2022).

32. N. T. Johansen, M. C. Pedersen, L. Porcar, A. Martel, L. Arleth, Introducing SEC–SANS for studies of complex self-organized biological systems. Acta Cryst D 74, 1178–1191 (2018).

33. S. R. Midtgaard, et al., Invisible detergents for structure determination of membrane proteins by small-angle neutron scattering. The FEBS journal 285, 357–371 (2018).

34. A. Raasakka, et al., Molecular structure and function of myelin protein P0 in membrane stacking. Sci Rep 9, 642 (2019).

35. S. Ruskamo, et al., Cryo-EM, X-ray diffraction, and atomistic simulations reveal determinants for the formation of a supramolecular myelin-like proteolipid lattice. J. Biol. Chem. 295, 8692–8705 (2020).

36. S. Ruskamo, et al., Atomic resolution view into the structure-function relationships of the human myelin peripheral membrane protein P2. Acta crystallographica.Section D, Biological crystallography 70, 165–176 (2014).

37. H. Fernández-morán, J. B. Finean, ELECTRON MICROSCOPE AND LOW-ANGLE X-RAY DIFFRACTION STUDIES OF THE NERVE MYELIN SHEATH. The Journal of Biophysical and Biochemical Cytology 3, 725–748 (1957).

38. D. L. Caspar, D. A. Kirschner, Myelin membrane structure at 10 A resolution. Nat New Biol 231, 46–52 (1971).

39. M. F. Moody, X-ray Diffraction Pattern of Nerve Myelin: A Method for Determining the Phases. Science 142, 1173–1174 (1963).

40. S. Suresh, C. Wang, R. Nanekar, P. Kursula, J. M. Edwardson, Myelin basic protein and myelin protein 2 act synergistically to cause stacking of lipid bilayers. Biochemistry 49, 3456–3463 (2010).

41. C. Stadelmann, S. Timmler, A. Barrantes-Freer, M. Simons, Myelin in the Central Nervous System: Structure, Function, and Pathology. Physiological Reviews 99, 1381–1431 (2019).

42. K. Woodward, S. Malcolm, CNS myelination and PLP gene dosage. Pharmacogenomics 2, 263–272 (2001).

43. G. Daffu, J. Sohi, J. Kamholz, Proteolipid protein dimerization at cysteine 108: Implications for protein structure. Neuroscience research (2012) https://doi.org/10.1016/j.neures.2012.07.009.

44. E. Swanton, A. Holland, S. High, P. Woodman, Disease-associated mutations cause premature oligomerization of myelin proteolipid protein in the endoplasmic reticulum. Proc Natl Acad Sci U S A 102, 4342–4347 (2005).

45. J. P. Schlebach, et al., Reversible Folding of Human Peripheral Myelin Protein 22, a Tetraspan Membrane Protein. Biochemistry (2013) https://doi.org/10.1021/bi301635f.

46. J. Zhao, et al., Multiple claudin–claudin cis interfaces are required for tight junction strand formation and inherent flexibility. Commun Biol 1, 1–15 (2018).

47. D. Boison, H. Büssow, D. D’Urso, H. W. Müller, W. Stoffel, Adhesive properties of proteolipid protein are responsible for the compaction of CNS myelin sheaths. J Neurosci 15, 5502–5513 (1995).

48. H. Inouye, H. Tsuruta, J. Sedzik, K. Uyemura, D. A. Kirschner, Tetrameric assembly of full-sequence protein zero myelin glycoprotein by synchrotron x-ray scattering. Biophys J 76, 423–437 (1999).

49. G. Krause, J. Protze, J. Piontek, Assembly and function of claudins: Structure-function relationships based on homology models and crystal structures. Semin Cell Dev Biol 42, 3–12 (2015).

50. J. Piontek, S. M. Krug, J. Protze, G. Krause, M. Fromm, Molecular architecture and assembly of the tight junction backbone. Biochimica et Biophysica Acta (BBA) - Biomembranes 1862, 183279 (2020).

51. L. Shapiro, J. P. Doyle, P. Hensley, D. R. Colman, W. A. Hendrickson, Crystal structure of the extracellular domain from P0, the major structural protein of peripheral nerve myelin. Neuron 17, 435–449 (1996).

52. H. Suzuki, K. Tani, Y. Fujiyoshi, Crystal structures of claudins: insights into their intermolecular interactions. Annals of the New York Academy of Sciences 1397, 25–34 (2017).

53. A. J. Thompson, M. S. Cronin, D. A. Kirschner, Myelin protein zero exists as dimers and tetramers in native membranes of Xenopus laevis peripheral nerve. J Neurosci Res 67, 766–771 (2002).

54. L. Montani, Lipids in regulating oligodendrocyte structure and function. Seminars in Cell & Developmental Biology 112, 114–122 (2021).

55. O. Spörkel, T. Uschkureit, H. Büssow, W. Stoffel, Oligodendrocytes expressing exclusively the DM20 isoform of the proteolipid protein gene: myelination and development. Glia 37, 19–30 (2002).

56. J. Tuusa, A. Raasakka, S. Ruskamo, P. Kursula, Myelin-derived and putative molecular mimic peptides share structural properties in aqueous and membrane-like environments. Multiple Sclerosis and Demyelinating Disorders 2, 4 (2017).

57. A. Denic, et al., The relevance of animal models in multiple sclerosis research. Pathophysiology 18, 21–29 (2011).

58. A. M. Messier, O. A. Bizzozero, Conserved fatty acid composition of proteolipid protein during brain development and in myelin subfractions. Neurochem Res 25, 449–455 (2000).

59. F. G. Mastronardi, C. A. Ackerley, L. Arsenault, B. I. Roots, M. A. Moscarello, Demyelination in a transgenic mouse: a model for multiple sclerosis. J Neurosci Res 36, 315–324 (1993).

60. M. E. de Bellard, Myelin in Cartilaginous Fish. Brain Res 1641, 34–42 (2016).

61. F. Schliess, W. Stoffel, Evolution of the myelin integral membrane proteins of the central nervous system. Biol Chem Hoppe Seyler 372, 865–874 (1991).

62. K. F. Mittendorf, et al., Peripheral myelin protein 22 alters membrane architecture. Science Advances (2017) https://doi.org/10.1126/sciadv.1700220 (February 8, 2022).

63. A. S. Dhaunchak, D. R. Colman, K.-A. Nave, Misalignment of PLP/DM20 Transmembrane Domains Determines Protein Misfolding in Pelizaeus–Merzbacher Disease. J. Neurosci. 31, 14961–14971 (2011).

64. N. Palaniyar, J. L. Semotok, D. D. Wood, M. A. Moscarello, G. Harauz, Human proteolipid protein (PLP) mediates winding and adhesion of phospholipid membranes but prevents their fusion. Biochim Biophys Acta 1415, 85–100 (1998).

65. M. Simons, E. M. Krämer, C. Thiele, W. Stoffel, J. Trotter, Assembly of myelin by association of proteolipid protein with cholesterol- and galactosylceramide-rich membrane domains. J Cell Biol 151, 143–154 (2000).

66. E. S. Mathews, et al., Mutation of 3-hydroxy-3-methylglutaryl CoA synthase I reveals requirements for isoprenoid and cholesterol synthesis in oligodendrocyte migration arrest, axon wrapping, and myelin gene expression. J Neurosci 34, 3402–3412 (2014).

67. E. S. Mathews, B. Appel, Cholesterol Biosynthesis Supports Myelin Gene Expression and Axon Ensheathment through Modulation of P13K/Akt/mTor Signaling. J Neurosci 36, 7628–7639 (2016).

68. A. E. Blaurock, D. L. Caspar, F. O. Schmitt, X-ray diffraction of living nerve. Small-angle diffraction. Neurosci Res Program Bull 9, 512–515 (1971).

69. A. I. Boullerne, The history of myelin. Exp Neurol 283, 431–445 (2016).

70. J. B. Finean, X-Ray Analysis of the Structure of Peripheral Nerve Myelin. Nature 173, 549–550 (1954).

71. D. A. Kirschner, D. L. Caspar, Comparative diffraction studies on myelin membranes. Ann N Y Acad Sci 195, 309–320 (1972).

72. G. Shahane, W. Ding, M. Palaiokostas, M. Orsi, Physical properties of model biological lipid bilayers: insights from all-atom molecular dynamics simulations. J Mol Model 25, 76 (2019).

73. A. E. Blaurock, C. R. Worthington, Low-angle X-ray diffraction patterns from a variety of myelinated nerves (1969) https://doi.org/10/2/60082ab6590a435ec9800d729061dbcd (February 10, 2022).

74. R. J. Chandross, J. B. Adams, R. S. Bear, Discontinuous transition of myelin structure at the junction between central and peripheral components of the eighth cranial nerve, as disclosed by X-ray diffraction. J Comp Neurol 177, 11–15 (1978).

75. P. Dupouey, et al., Immunochemical studies of myelin basic protein in shiverer mouse devoid of major dense line of myelin. Neurosci Lett 12, 113–118 (1979).

76. A. Privat, C. Jacque, J. M. Bourre, P. Dupouey, N. Baumann, Absence of the major dense line in myelin of the mutant mouse “shiverer.” Neurosci Lett 12, 107–112 (1979).

77. A. Thureau, P. Roblin, J. Pérez, BioSAXS on the SWING beamline at Synchrotron SOLEIL. J Appl Cryst 54, 1698–1710 (2021).

78. C. E. Blanchet, et al., Versatile sample environments and automation for biological solution X-ray scattering experiments at the P12 beamline (PETRA III, DESY). J Appl Cryst 48, 431–443 (2015).

79. K. Manalastas-Cantos, et al., ATSAS 3.0: expanded functionality and new tools for small-angle scattering data analysis. J Appl Cryst 54, 343–355 (2021).

80. P. Asthana, et al., Structural insights into the substrate-binding proteins Mce1A and Mce4A from Mycobacterium tuberculosis. IUCrJ 8, 757–774 (2021).

81. A. Calcutta, et al., Mapping of unfolding states of integral helical membrane proteins by GPS-NMR and scattering techniques: TFE-induced unfolding of KcsA in DDM surfactant. Biochim Biophys Acta 1818, 2290–2301 (2012).

82. J. Døvling Kaspersen, et al., Low-resolution structures of OmpA.DDM protein-detergent complexes. Chembiochem 15, 2113–2124 (2014).

83. S. L. Harwood, et al., Structural Investigations of Human A2M Identify a Hollow Native Conformation That Underlies Its Distinctive Protease-Trapping Mechanism. Molecular & Cellular Proteomics 20, 100090 (2021).

84. D. I. Svergun, et al., Protein hydration in solution: Experimental observation by x-ray and neutron scattering. PNAS 95, 2267–2272 (1998).

85. D. I. Svergun, M. V. Petoukhov, M. H. Koch, Determination of domain structure of proteins from X-ray solution scattering. Biophysical journal 80, 2946–2953 (2001).

